# Long-Term Functional 3D Human Skeletal Muscle Constructs from Human Induced Pluripotent Stem Cells Enable Electrically Controlled Remodelling Studies

**DOI:** 10.1101/2025.08.28.672872

**Authors:** C. Rivares Benitez, N.E.M. Klein Hofmeijer, E. Feola, J. Rouwkema, M. Sartori

## Abstract

Studying skeletal muscle remodelling across large temporal scales is difficult because most 3D tissue-engineered skeletal muscle (3D-TESM) models cannot sustain stable development over prolonged culture. Skeletal muscle adaptations in vivo are strongly influenced by changes in tissue length, but most TESM models are mechanically restrictive, limiting elongation and thereby constraining structural remodelling. The stiffness of the pillars that anchor the tissues is a key determinant of how much they can shorten or elongate during development, thereby shaping both their morphology and force-generating capacity. To address this limitation, we investigated how TESMs derived from human induced pluripotent stem cells (hiPSCs) develop over two weeks when suspended between pillars with distinct mechanical properties, ranging from mechanically compliant to mechanically stiff boundary conditions. We focused on both passive and active force-generating properties.

With compliant pillars, tissues compacted primarily along their length (~66% reduction), followed by progressive elongation (~50% increase) from day 5 onward. In contrast, with stiff pillars, tissues compacted across their width, with no longitudinal shortening. Despite these morphological differences, projected tissue area decreased similarly across setups. Spontaneous contractions emerged from day 7 and continued until day 14, typically occurring in bursts at 0.6–0.8 Hz. Passive force peaked early with compliant pillars and declined thereafter, while remaining relatively stable with stiff pillars. Under stepwise electrical stimulation, maximal contractility was consistently observed at ~50 Hz. Active force rose sharply between day 12 and day 14, with the steepest increase (up to 2000%) in compliant pillars. Although sample size was limited, the consistency of trends across stiffness conditions and surface treatments provides valuable insights into how boundary mechanics shape 3D-TESM development. These findings offer a foundation for designing next-generation muscle culture systems that support dynamic remodelling, force maturation, and training-responsive behaviours.

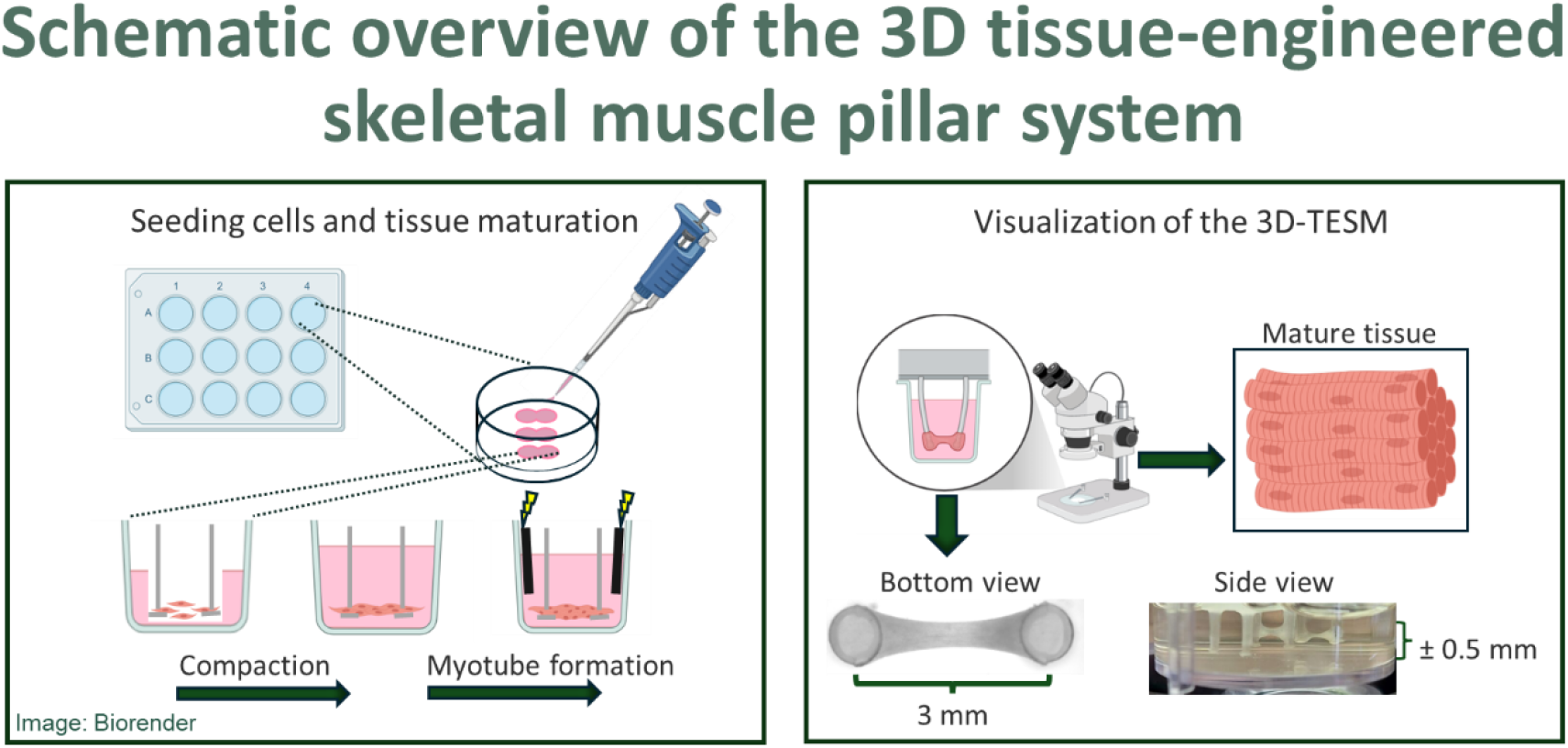

## Introduction

Skeletal muscles are highly plastic^1,2^. Muscle fibres can adapt their architecture by adding or removing sarcomeres in series, affecting fibre length, or in parallel, modifying force-generating capacity^2,3^. Other key properties, such as endurance capacity (i.e. the ability to shift fibre phenotypes), also change over time in response to physiological or mechanical stimuli^4,5^. Accurately guiding muscle plasticity and remodelling using external electro-mechanical cues could significantly improve rehabilitation strategies and tailored training interventions for athletes. However, there are still fundamental knowledge gaps in understanding how human skeletal muscles remodel in response to electro-mechanical stimuli. This gap is particularly critical at the extremes of temporal scales, across weeks and months, where most physiologically relevant adaptations occur.

In vivo studies have provided valuable insight, but they remain complicated by the difficulty of isolating the effect of a single external stimulus on muscle remodelling. Muscles in the body are physically connected to synergists and antagonists through connective tissue, are innervated, and are constantly exposed to systemic cues, all of which confound interpretation of structural adaptation to a defined input.

In vitro approaches offer a promising solution by enabling precise control of the environment. However, most current 3D-TESM constructs rely on passive systems, where the fixed spacing between pillars mechanically constrains tissue elongation^6–10^. This is a significant limitation, as many physiologically relevant adaptations involve changes in tissue length, including natural muscle growth during development, velocity- and power-specific strength training, and clinical interventions such as lengthening overly shortened muscles in patients with neuromuscular disorders (e.g., spastic cerebral palsy). Moreover, few studies have examined human skeletal muscle in vitro over extended timescales^6–10^, and even fewer have addressed how mechanical and electrical cues interact to shape remodelling across multiple weeks. Sustaining tissue health and function for long-term analyses remains a key challenge.

To begin addressing this, we conducted an exploratory study to examine how engineered muscle tissues respond over time when suspended between compliant (low-stiffness) and stiff (high-stiffness) pillars. Pillar compliance directly affects the magnitude of mechanical stimuli experienced by the tissues: compliant pillars allow greater shortening and elongation, whereas stiff pillars restrict movement and impose higher resistance. We modified an existing in vitro platform^6^ that supports the formation of human 3D-TESM in a controlled environment and allows precise manipulation of mechanical, biochemical, and structural cues. Importantly, these tissues contain myogenic progenitor cells capable of sarcomerogenesis^11^, providing a model to study how sub-cellular and tissue-scale adaptation relates to specific training-like interventions. In this study, we focus on the effects of electrical pacing over two weeks, highlighting the potential of this system to support analyses of mid- to long-term remodelling. This novelty directly relates to rehabilitation following stroke or cerebral palsy, as well as to tailored training interventions for athletes aiming to improve power generation and contraction speed.

## Methods / experimental section

### Fabrication of platform

Tissue culture platforms, similar to that described by Ribeiro et al. (2022)^6^, were used in this study. The base consisted of a standard commercial 12-well plate (Greiner Bio-One, Netherlands), into which custom moulds with tissue slots were inserted. Each mould was designed to fit within a well and contained three small tissue slots for cell seeding. Importantly, the slots were identical in size across all mould types, ensuring that only the mould material differed. Three types of moulds were used to form the tissue slots:

Gelatine moulds were prepared by dissolving gelatine (from porcine skin) powder (8.2% w/v, Sigma-Aldrich) in Milli-Q water at 60 °C. The solution was autoclaved, DMEM (55.6% v/v, Gibco) was added. At 37 °C, 1.7 mL of the gelatine solution was pipetted into each well of a 12-well plate. A negative mould, shaped using tissue-specific templates, was inserted to define the geometry of the tissue slots. The plates were placed at 4 °C for 4 hours and then frozen overnight to ensure solidification. Once fully gelled, the moulds were carefully removed, leaving behind defined slots suitable for cell seeding.

PDMS moulds were cast using the same tissue-shaped templates as above mentioned. The PDMS was cured for 12 hours at 65 °C, after which the moulds were treated with Pluronic F-127 (1% Pluronic F-127 in PBS) to reduce cell adhesion. They were then sterilized by immersion in 70% ethanol for 15 minutes, rinsed three times with PBS, and exposed to UV light for two 30-minute cycles.

Polyamide (PA2200) moulds were 3D printed using a SLS printer (FORMIGA P101, EOS GmbH) with the same slot geometry. PA2200 was chosen for its cytocompatibility with iPSC-derived cells and its ability to withstand autoclaving^12^. These rigid moulds offered several practical advantages over gelatine-based ones, which liquefy at 37 °C and compromise tissue support. Compared to PDMS, PA2200 moulds were easier to produce and did not require surface treatments such as Pluronic F-127, which causes hydrophobic surface. However, a limitation of the PA2200 moulds is that they block visual access to the tissues using an inverted microscope.

PDMS Pillars were fabricated in-house using PDMS at a 1:10 curing agent to base ratio. The PDMS was cast into pillar-shaped moulds, cured for 12 hours at 65°C, and sterilized following the same protocol as the PDMS moulds.

Thermoplastic elastomer (TPE) pillars kindly provided by prof. Passier’s group from the University of Twente. The TPE pillars exhibit significantly higher stiffness compared to PDMS pillars, as previously characterized by Rivera-Arbeláez et al. (2023)^13^. See Figure 1 for a graphical overview of the different platform configurations.

**Figure 1.**
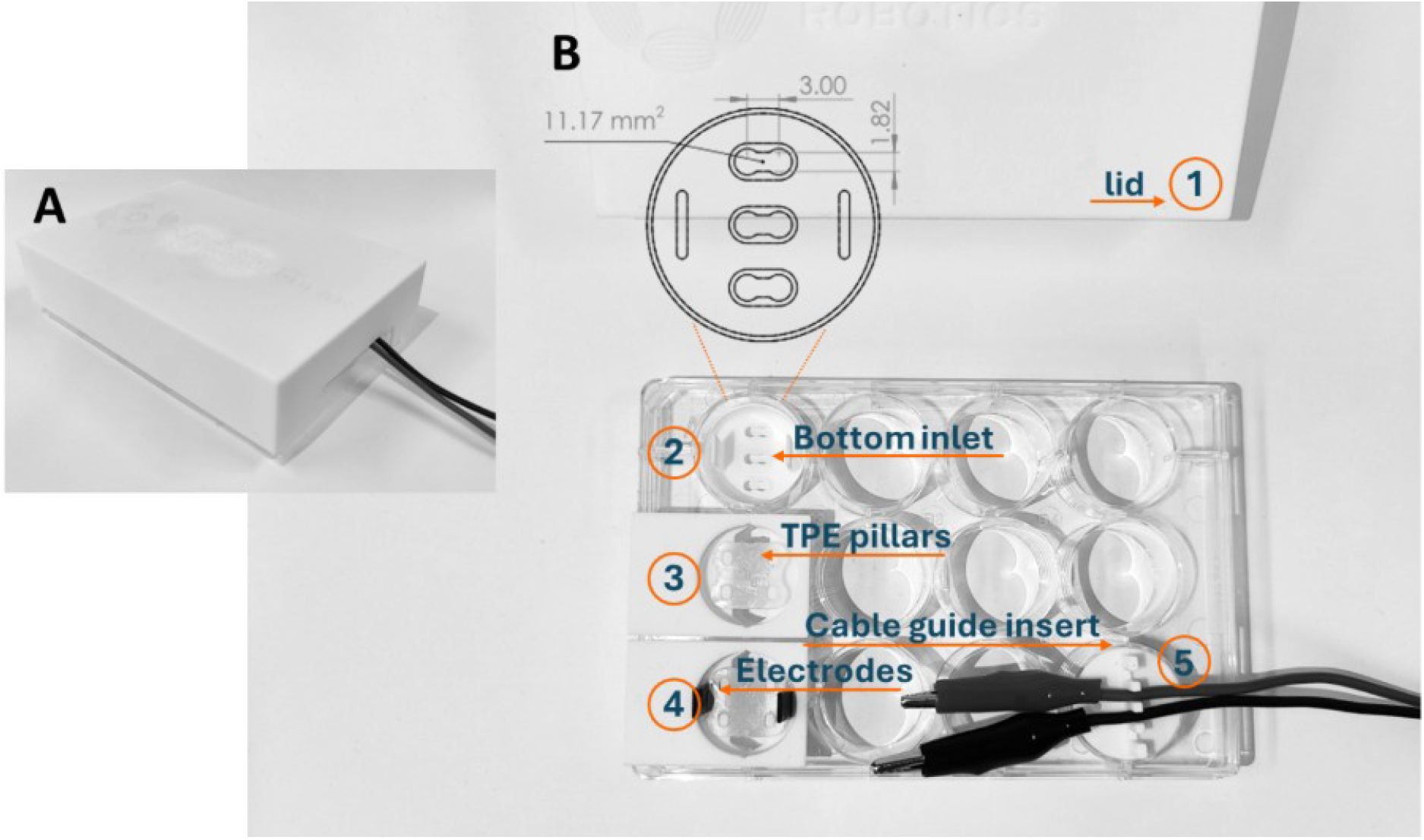
Overview of the culture system. Inspired by the “Versatile System Platform” (Ribeiro et al. 2022)^6^, we modelled custom components for a 12-well plate with a modified lid to support tissue engineering workflows (Figure 1A-B). The design includes a lid (*1) with an entry slot to allow cable access for controlled tissue pacing (Figure 1A-B). Specifically, well 1 includes a mould inlet (*2) for precise cell pipetting and tissue moulding based on the calculations described in Ribeiro et al. (2022)^6^. Well 5 contains a pillar holder with TPE pillars (*3) to guide tissue formation, and well 9 integrates electrodes for electrical stimulation (*4) and functional assessment. Additionally, well 12 houses a cable guide insert (*5).

**Figure 2.**
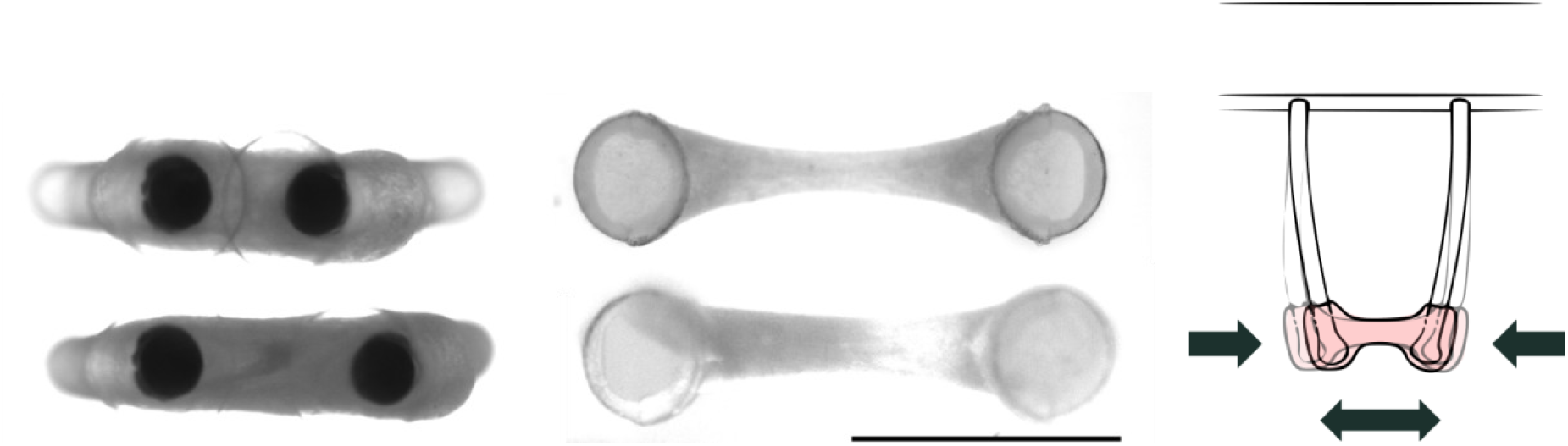
Representative images of two typical 3D-TESMs cultured on PDMS and TPE pillars at day 5 and day 15. 3D-TESMs on PDMS pillars exhibit noticeable compaction over the first 5 days post-differentiation, as indicated by a decrease in tissue length. In contrast, 3D-TESMs on TPE pillars primarily show a reduction in tissue width during the first 5 days post-differentiation, with minimal changes in length. The scale bar represents 2000 µm.

### Formation of 3D-TESM of myogenic progenitors

Myogenic progenitor cells were kindly donated by prof. Pijnappel from the Erasmus MC in Rotterdam (the Netherlands). A transgene-free method was used by the previously named group for the generation of purified, expandable myogenic progenitors from hIPSc, as mentioned in ref: van der Wal 2017^14,15^. Myogenic progenitor cells were expanded in high-glucose DMEM (Gibco, Waltham, MA) supplemented with 10% fetal bovine serum (Hyclone, Thermo Scientific, Waltham, MA), 1% penicillin-streptomycin-glutamine (Gibco, Waltham, MA), and 100 ng·mL^-1^ FGF2 (Prepotech, Rocky Hill, NJ), on extracellular matrix-coated-coated dishes (1:200, Sigma-Aldrich, E6909). The medium was refreshed every other day. For passaging, cells were detached with TrypLE reagent (Gibco, Waltham, MA). For cryopreservation, cell pellets were resuspended in proliferation medium with 10% DMSO (Sigma-Aldrich), using standard freeze-thaw procedures.

3D-TESMs were engineered by mixing the myogenic progenitors (250.000) resuspended in DMEM with a hydrogel formulation that contained 10% v/v fibrinogen (dissolved in DMEM, final concentration 2 mg·mL^−1^, Sigma-Aldrich), 20% v/v Matrigel Growth Factors-Reduced (Corning), 69% v/v proliferation medium. After well mixing of the hydrogel mix and MP, 1% v/v Thrombin (dissolved in 0.1% BSA, final concentration 20 U·mL^−1^, Sigma-Aldrich) was mixed into the mix. The hydrogel formulation was based on thepaper of Iuliano et al. (2023)^4^. A total volume of 15 uL of the cell-hydrogel mix was pipet into each tissue slot within the 12-well plates. After pipetting of the 3 tissue slots the holder plus pilar system, which were previously assembled, were immediately placed in the well. The tissues were left for 10 minutes in the laminar airflow cabinet, whereafter 1ml of proliferation medium was added to each well and the system was placed into the incubator (set at 37°C and 5% CO2). 1mL of the Medium was changed after 24 hours and After 48 h (D0), the proliferation medium was replaced by differentiation medium consistent of DMEM with high (4.5 g·mL^−1^) glucose, 1% v/v 1% penicillin-G (Sigma-Aldrich), 1% v/v KnockOut Serum Replacement (ref), 1% v/v GlutaMAX supplement (Gibco), 1% v/v ITS-X Insulin-Transferrin-Selenium-Ethanolamine (Gibco) and 1.5 mg·mL^−1^ 6-aminocaproic acid (Sigma-Aldrich). After 3 days (D3), half the volume of differentiation medium was replaced by differentiation medium enriched to 20% v/v KO. Half of the medium was refreshed every other day until electrical stimulation protocol was started.

### Electrical Stimulation

The 3D-TESMs were electrically stimulated using 2 carbon fibre electrodes (3 x 1 mm carbon fibre strip, EasyComposites, EU), spaced 16 mm apart. Electrical pulse stimulation was performed by a stimulus generator (STG5, Multichannel systems, Germany), this system was used for training as well as force measurements. The stimulations were current controlled and had a square bipolar waveform. To assess force generation, tissues were subjected to a stepwise electrical stimulation protocol. First, stimulation amplitude was increased from 5 mAto 15 mAin 5 mA increments. At each amplitude level, pulse width was varied sequentially from 20 ms to 3 ms. Specifically, 20 pulses were applied for 20 ms and 10 ms pulse widths, and 70 pulses for 5 ms and 3 ms pulse widths.

For contractile training, tissues were stimulated similar to that of the characterization protocol. A total of 50 pulses were delivered at 8 mA amplitude (square bipolar waveform) with 5 ms pulse width.

### Live imaging and imaging analysis

Tissues were imaged using an Evident IX83 inverted wide-field fluorescence microscope (Evident, Japan) equipped with a 2× objective. Pillar movements were quantified using Tracker software (version 6.3.1; Brown, 2009) to obtain exact pillar displacement and Muscle Motion^16^ plugin for ImageJ (version 1.54p, 17 February 2025; Schneider et al., 2012) to assess pixel-level motion. Morphological features were extracted using a custom ImageJ macro, while a custom-built Python analysis pipeline (https://github.com/CEINMS-RT/invitro_morphological_analysis) was tested and is currently under development.

In the Python pipeline, pillar locations were identified using a Hough transform-based detection algorithm, with candidate centroids filtered to retain only the two supporting pillars. The inter-pillar distance was converted to micrometers using microscope-specific calibration scales and applied to the elastic beam bending equation (Serrao et al., 2012). To isolate the tissue region from the background, a GrabCut segmentation procedure was applied, initialized by intensity thresholds and the detected pillar locations. The segmented tissue was exported as a mesh, and the extracted contour was used to estimate tissue area.

### Characterization of 3D-TESMs contractile properties

The passive and peak force produced by the 3D-TESM on PDMS and TPE pillars over time were calculated by the elastic beam bending equation (Serrao et al. 2012). Where Young’s modulus was to 1MPa (known to vary between 0.5 and 3.5 MPa in between batches), the radius of the PDMS pilar and the length of the pilar were 0.25 mm and 3 mm, respectively. The height of the 3D-TESM on the pillar was not exactly known, but assuming from previous pilots the tissue width at close to the pillars, we assume 2.5 mm from the base of the pillars. Morphological and passive force data were fitted using second-order polynomial (quadratic) least-squares regression.

Graphs were generated using GraphPad Prism (GraphPad Software v.10.4.2, San Diego, CA).

## Results

### Early tissue compaction and morphological changes in 3D-TESMs

#### 3D-TESMs compaction

Tissue formation occurred rapidly, with constructs forming around the pillars typically within 10 minutes after seeding. During the first 7 days of culture (Day-2 to Day5), tissues underwent substantial compaction (~66%), as indicated by a reduction in inter-pillar distance (Fig. 3A). This early compaction phase was consistent across all mould types, gelatine, PA2200, and PDMS, which showed comparable reductions in tissue length along the pillar axis. In contrast, tissues formed around TPE pillars exhibited a markedly different pattern (Fig. 3A). Although a similar degree of compaction (~60%) was observed during the same early period, this occurred primarily across the tissue width rather than its length (Fig. 3B).

**Figure 3.**
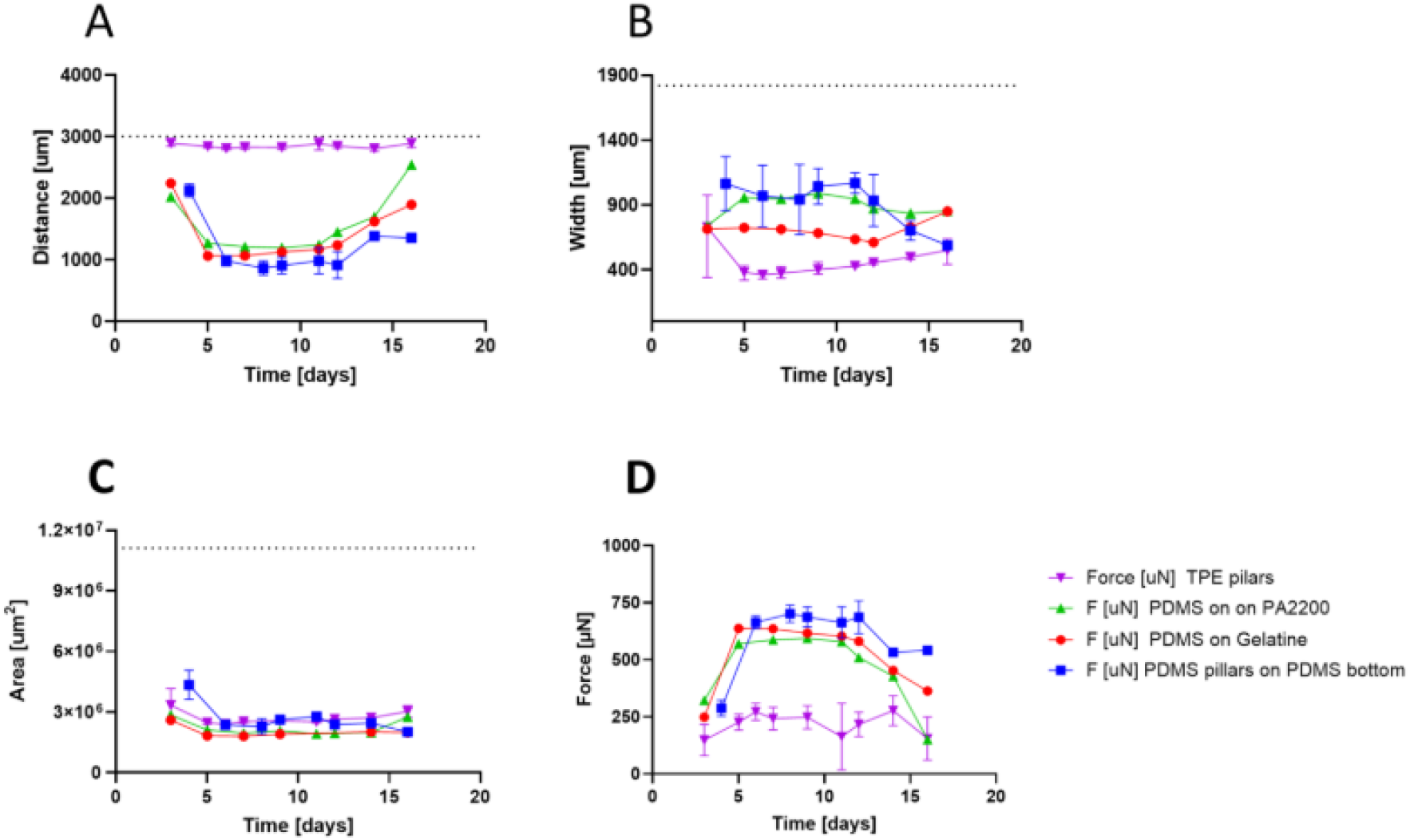
Morphological development and passive force in 3D-TESM over 2 weeks. **A)** Inter-pillar distance decreased rapidly during the first 5 days post-differentiation for 3D-TESMs on PDMS pillars, indicating early compaction. In contrast, TPE-based constructs maintained stable length. After compaction, PDMS-based tissues elongated exponentially, approaching the maximum spacing (~3 mm). **B)** Tissue width at the pillar midpoint decreased in TPE-based constructs during compaction, then gradually increased; PDMS-based tissues showed the opposite trend. **C)** Projected tissue area remained similar across conditions, suggesting that total compaction was comparable, with thedirection (length vs. width) influenced by pillar stiffness. **D)** Passive force (from pillar deflection) increased sharply during early compaction on PDMS pillars, then declined as tissues elongated. On TPE pillars, passive force could not be accurately quantified due to minimal deflection. Dashed lines indicate initial values at seeding. Note, we will increase the distance of the TPE pillars by 0.2mm, so figures make more visual sense

After day 5 post-differentiation, tissue morphology on PDMS pillars shifted notably, with constructs progressively elongating along the pillar axis (Fig. 3A). In contrast, tissues on TPE pillars began to gradually increase in width over the same period (Fig. 3B).

Despite the distinct compaction patterns observed on PDMS and TPE pillars, the projected tissue area (as viewed from below) decreased to a similar extent across all pillar types (Fig. 3C).

#### Morphological changes in 3D-TESMs after initial compaction

Following the initial compaction phase, 3D-TESM on PDMS pillars displayed marked elongation beginning around day 5 post-differentiation, with tissue length increasing along an apparent exponential trajectory (Fig. 3A). This elongation can be described by the equation: Length = 537 µm × exp (0.07 × days), with a tissue length doubling time of 9.2 days (R^2^ = 0.72).

In contrast, tissues formed on TPE pillars began to gradually increase in width around day 5 post-differentiation, following a similar exponential trend described by: Width = 289 µm× exp (0.04 × days), with a tissue width doubling time of 18 days (R^2^ = 0.68; Fig. 3B).

### Functional characteristics

#### Passive force generated by 3D-TESM over time

During the initial compaction phase, 3D-TESMs on PDMS pillars exerted increasing force on the pillars, as evidenced by progressive inward deflection and a corresponding decrease in inter-pillar distance. This phase was followed by a short plateau period, after which the passive force gradually declined over time. This decline can be described by the equation: Passive force = 450 + 53.0 × days – 3.5 × days^2^ (R^2^ = 0.79; Fig. 3D).

In contrast, tissues formed on TPE pillars did not follow this trend. Instead, the passive force remained relatively stable throughout the culture period, showing no clear compaction-driven increase or subsequent decline of force (Fig. 3D).

#### Emerging of spontaneous contractions

Spontaneous contractions were first observed in 3D-TESM cultured on both PDMS and TPE pillars starting around day 7 post-differentiation, see figure 4 for typical example of these contractions. The spontaneous contractions lasted in this manner till day 14. The contractions occurred without external stimulation and were noted to become more frequent following medium changes and electrical pacing. The frequency of spontaneous contractions varied across samples but typically ranged between 0.6 and 0.8 Hz. Although not strictly rhythmic, contractions often occurred in bursts.

**Figure 4.**
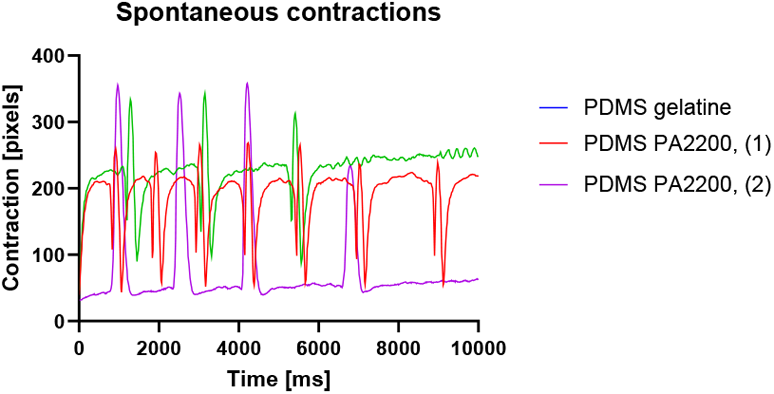
Spontaneous contractions in 3D-TESM. Representative examples of a single 3D-TESM exhibiting spontaneous contractions at three time points within one day. Bursts typically occur every few minutes between days 7 and 14 post-differentiation. After the second week, spontaneous activity becomes less frequent but remains detectable, particularly following electrical stimulation. Within bursts, contraction frequency begins around 0.6-0.8 Hz and tapers off after a few seconds. Timing and frequency are variable and appear random. Spontaneous contractions often intensify after medium changes and are observed across all 3D-TESMs, regardless of material.

After day 14 the spontaneous were typically present after applying electrical pulse stimulation (e.g. Fig. 5C-D).

**Figure 5.**
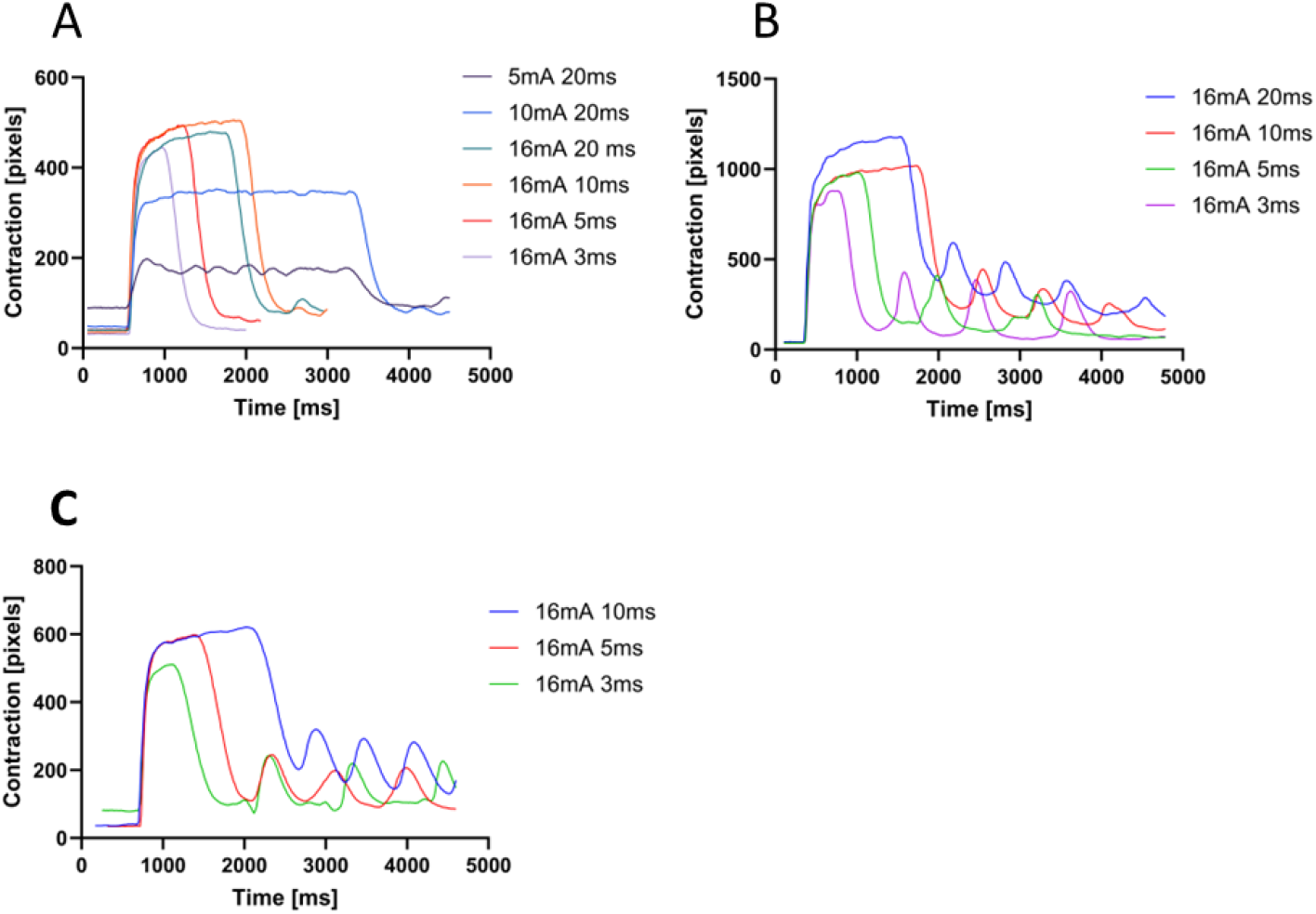
Contractile responses of 3D-TESM upon electrical stimulation at day 12 post-differentiation. Shown are representative contraction traces from individual 3D-TESMs subjected to a stepwise stimulation protocol with increasing pulse amplitudes (5 mA to 16 mA) and frequencies (25, 50, 100, and 167 Hz). As shown in panel A, stimulation amplitude was first increased to ensure maximal contraction, consistent with physiological recruitment. Once the maximal response was achieved, stimulation frequency was progressively increased. Peak contractions typically occurred around 50 Hz using 70 bipolar pulses of 10 ms each; higher frequencies did not further enhance contractile output. **A)** Example from TPE pillars with PA2200 mould. **B)** Example from PDMS pillars with PA2200 mould. **C)** Example from PDMS pillars with gelatine mould.

#### Contractility assessment via electrical stimulation (contraction dynamics)

During stepwise electrical stimulation, maximal contractions were consistently elicited using pulse trains of approximately 70 bipolar pulses, with pulse durations of 20, 10, 5, and 3 ms, corresponding to stimulation frequencies of 25, 50, 100, and 167 Hz, respectively. Increasing the stimulation frequency beyond this range did not enhance contractile force (see Fig. 5A for a representative example). The highest contractility was observed at a pulse duration of approximately 10 ms (50 Hz), with similar results obtained in both PDMS and TPE pillar systems (Fig. 5A-D).

#### Active force assessment

Active tissue force increased substantially between day 12 and day 14 post-differentiation across tissues cultured on two distinct pillar materials (PDMS and TPE).

Tissues formed on TPE pillars exhibited a modest but consistent increase in pillar deflection on day 14 compared to day 12, across all stimulation frequencies. A representative example is shown in Figure 6A. Correspondingly, active force increased by 304% to 433%, measured at 50, 100, and 167 Hz.

**Figure 6.**
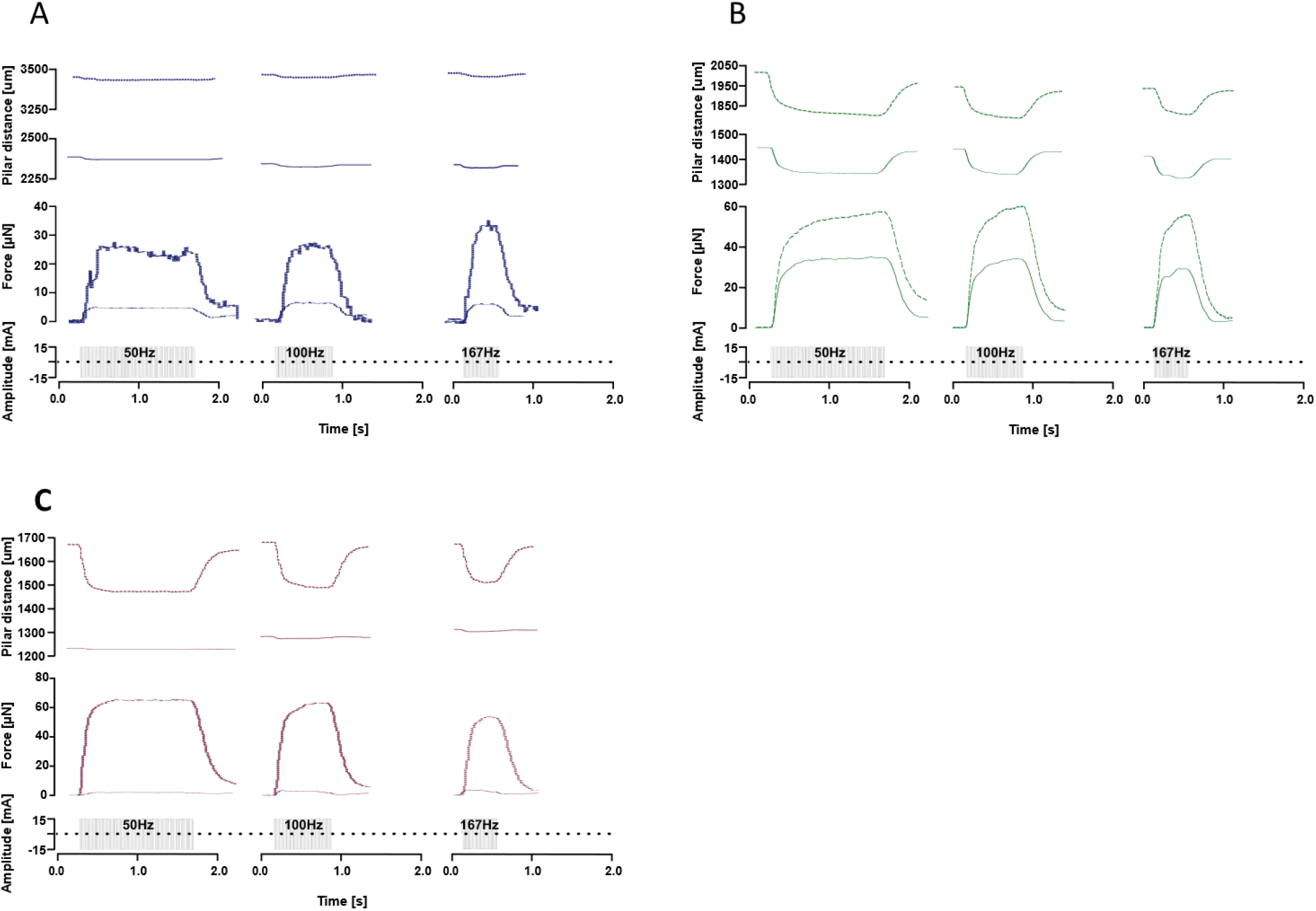
Active force exertion on pillars and pillar displacement under different stimulation frequencies at 12- and 14-days post-differentiation. **A)** 3D-TESMs on TPE pillars moulded with PA2200 at day 12 and day 14. **B)** 3D-TESMs on PDMS pillars using a gelatine-coated mould at day 12 and day 14. **C)** 3D-TESMs on PDMS pillars moulded with PA2200 at day 12 and day 14. Each subfigure shows pillar displacement at three stimulation frequencies (50, 100, and 166 Hz). Solid lines represent force and pillar displacement at day 12, and dashed lines represent day 14.

The most pronounced increase in both pillar deflection and activeforcewas observed in tissues cultured on PDMS pillars with a gelatine-coated bottom, where active force increased ranging from 1460% to 2000% between day 12 and day 14 across all stimulation frequencies (Figs. 6B).

Tissues on PDMS pillars with a PA2200 mould also showed substantial increases in deflection and active force, ranging from 48% to 65% over the same time period and frequency range (Figs. 6C).

In contrast, tissues formed on PDMS pillars with a PDMS mould exhibited minimal deflection, preventing accurate quantification of active force.

Among all tested conditions, PDMS pillars with a gelatine-coated mould yielded not only the largest increase in active force between day 12 and day 14 (mean = 29.1 ± 3.4 µN / 2 days) (Fig.7), but also the highest absolute force values on day 14. Tissues cultured on PDMS pillars with a PA2200 mould generated slightly lower active forces at that time point and exhibited a correspondingly lower slope of force increase (mean = 8.89 ± 2.16 µN / 2 days). Finally, tissues on TPE pillars showed a slightly higher slope of active force increase compared to those on PDMS pillars with PA2200 bottoms (mean = 12.01 ± 2.1 µN / 2 days).

**Figure 7.**
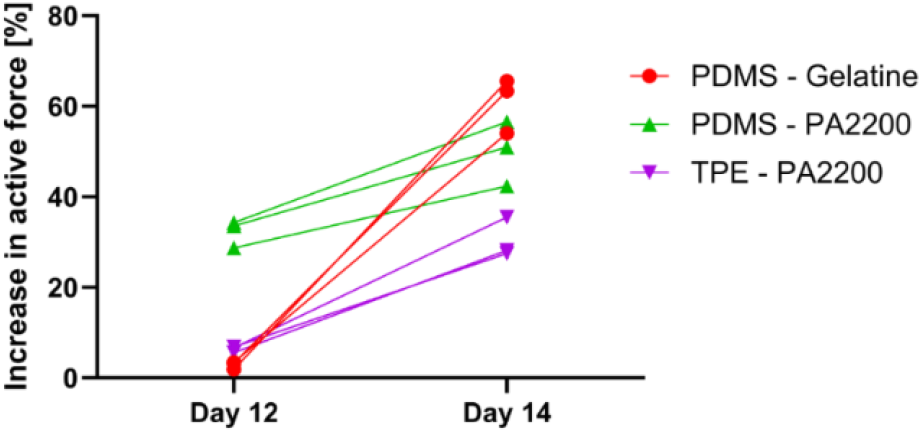
Active Force Progression at 12- and 14-days post-differentiation. This figure shows the percent increase in active tissue force from day 12 to day 14 post-differentiation across different culture conditions. Tissues were formed on either PDMS or TPE pillars, combined with gelatine- or PA2200-mould surfaces. Each line represents stimulation frequency.

## Discussion/Implications

In this study, we developed a culture system that sustains human 3D-TESM under electrical pacing for up to two weeks. By varying pillar stiffness and mould conditions, the platform enables tissues to compact, elongate, and progressively acquire contractile function, providing a framework to study remodelling across mid- to long-term timescales. This system provides a framework for studying muscle remodelling across mid- to long-term timescales and represents a first step toward modelling training-like interventions in vitro, where repeated electrical and mechanical inputs drive structural and functional adaptations.

### Transition from compaction to growth phase

The compaction phase is a critical yet often underreported step in 3D-TESM development^17^. During this phase, muscle progenitor cells generate contractile forces on the hydrogel matrix, producing tissue shrinkage and alignment between anchoring structures. This reorganization induces passive tension that facilitates alignment, myotube fusion, and early maturation^17–19^. In our study, compaction proceeded for ~7 days, spanning the proliferation and early differentiation phases. The direction of compaction was determined by pillar mechanics, occurring lengthwise on PDMS pillars versus across the width on TPE pillars. Notably, tissues on compliant PDMS pillars exhibited the highest active force after two weeks, reinforcing the functional importance of supporting compaction.

### Growth phase following compaction

After compaction, PDMS-anchored tissues entered a growth phase, elongating exponentially with a doubling time of ~9 days. Growth reflects a coupled process of hydrogel degradation and new ECM deposition^20^, which stretch the construct and alter compliance. Tissues with higher Matrigel concentrations are known to produce thicker myofibers and higher forces, supporting this interpretation. The observed decline in passive force aligns with scaffold turnover combined with tissue-driven elongation. In vivo, passive and active tension are key drivers of sarcomerogenesis ^21^; here, pillar-applied passive tension combined with spontaneous and evoked contractions likely underpinned the rapid length growth we observed.

### Functional maturation and stimulation response

Spontaneous contractions appeared around day 7 and persisted through day 14, increasing with medium changes and pacing. By day 14, the strongest contractions occurred at ~50 Hz stimulation, with no further gains at higher frequencies. Active force rose sharply between day 12 and 14, with PDMS-gelatine conditions showing increases of up to 2000%. This dramatic rise suggests coordinated myotube maturation, ECM reinforcement, and efficient force transmission, a hallmark of a training-like adaptation in vitro.

### Repeatability and variability

Overall trends were consistent, but the repeatability of specific measurements varied between readouts. In particular, it is difficult to assess how consistent active force generation truly is across tissues, as substantial differences were observed between pillar materials and even between mould types within the same material. This high degree of variability is not unique to our work; several groups have also reported considerable variability in active force output between constructs^7^. By contrast, passive force trajectories and morphological changes over time were much more consistent, showing comparable compaction and growth dynamics across conditions. To our knowledge, however, no other studies have systematically tracked passive force or morphology longitudinally as a quality indicator, representing a current gap in the literature. Notably, PDMS-gelatine pillars yielded the strongest and most reproducible active force results in our hands. Longitudinal monitoring of passive force during compaction therefore emerged as a robust, non-invasive indicator of tissue health and may help flag suboptimal constructs early in culture.

### Limitations

A key limitation of this study is the low number of samples, which reduces the power of the findings. However, the observed trends were consistent across multiple time points and experimental setups, lending confidence to the biological relevance of the results. Despite the limited sample size, these findings provide valuable insights that can inform the design of next-generation 3D skeletal muscle culture systems, both in our own work and across the broader field.

### Future work and active lengthening

Future studies should expand sample numbers and scale up the system for parallel testing. A central objective is the development of a reliable and scalable culture system that enables systematic, longitudinal experimentation and accelerates progress toward functional 3D-TESM models for training-like interventions.

Longitudinal tracking of passive force, particularly during the compaction phase, could serve as a non-invasive readout of tissue health and remodelling state. By correlating force evolution with in vivo immunohistology and molecular benchmarks, it may be possible to define biologically relevant transitions that bridge early formation with later functional development. Likewise, spontaneous contractions, reflecting functional self-organization^17-19^, could be standardized and scored as an early indicator of readiness for electrical or mechanical stimulation, helping to prevent premature or excessive loading.

A critical gap in the field is the absence of systematic studies on longitudinal stimulation of engineered muscle tissues. To our knowledge, no work has compared different stimulation types over extended timescales, meaning that the long-term consequences of over- or under-stimulation remain unknown. Similarly, there are currently no systems capable of applying controlled active lengthening to 3D-TESM, let alone combining lengthening with electrical pacing. This represents a major limitation, as in vivo muscle training and rehabilitation rely on the interplay between loading and excitation.

Addressing these gaps will require adaptive, feedback-driven pacing schemes that adjust to tissue response over time, as well as new culture platforms capable of applying mechanical strain in combination with electrical input. Establishing such reproducible culture systems that support substantial longitudinal tissue lengthening in response to training-like stimuli should be a central objective for the community. Our present study, while focused on electrical pacing, provides a foundation for developing and testing these next-generation protocols, directly modelling the key drivers of adaptation in both rehabilitation and athletic performance.

## Conclusion

This study shows the importance of the compaction phase for functional maturation in 3D-TESMs. We show that the mechanical properties of the culture platform strongly dictate tissue remodelling trajectories, influencing elongation and force development. These findings establish passive force evolution and tissue morphology as useful longitudinal indicators of construct health and adaptation. To advance toward physiologically relevant growth and training applications, next-generation systems should combine controlled length modulation with electrical stimulation in scalable and reproducible formats.

## Acknowledgements

The thank dr. José Manuel Rivera-Arbelaez for his guidance in the methods required for material preparation. We also acknowledge dr. Alessandro Iuliano and prof. Pim Pijnappel for providing the cells and for their valuable advice on initiating the cell cultures and optimizing the hydrogel composition.

## Funding statement

This work was supported in part by the European Research Council (ERC) under the European Union’s Horizon 2020 Research and Innovation Program, as part of the ERC Consolidator Grant ROBOREACTOR under Grant 101123866.

## Notes

### Competing Interest Statement

The authors have declared no competing interest.

### Summary of Updates

This version corrects an acknowledgment error: the source of the cells was misattributed and is now accurately stated in the Methods section ("Formation of 3D-TESM of myogenic progenitors"). No data, results, or conclusions were changed.

